# Interpretable, scalable, and transferrable functional projection of large-scale transcriptome data using constrained matrix decomposition

**DOI:** 10.1101/2021.04.13.439654

**Authors:** Nicholas Panchy, Kazuhide Watanabe, Tian Hong

## Abstract

Large-scale transcriptome data, such as single-cell RNA-sequencing data, have provided unprecedented resources for studying biological processes at the systems level. Numerous dimensionality reduction methods have been developed to visualize and analyze these transcriptome data. In addition, several existing methods allow inference of functional variations among samples using gene sets with known biological functions. However, it remains challenging to analyze transcriptomes with reduced dimensions that are interpretable in terms of dimensions’ directionalities, transferrable to new data, and directly expose the contribution of individual genes. In this study, we used gene set non-negative principal component analysis (gsPCA) and non-negative matrix factorization (gsNMF) to analyze large-scale transcriptome datasets. We found that these methods provide low-dimensional information about the progression of biological processes in a quantitative manner, and their performances are comparable to existing functional variation analysis methods in terms of distinguishing multiple cell states and samples from multiple conditions. Remarkably, upon training with a subset of data, these methods allow predictions of locations in the functional space using data from experimental conditions that are not exposed to the models. Specifically, our models predicted the extent of progression and reversion for cells in the epithelial-mesenchymal transition (EMT) continuum. These methods revealed conserved EMT program among multiple types of single cells and tumor samples. Finally, we demonstrate this approach is broadly applicable to data and gene sets beyond EMT and provide several recommendations on the choice between the two linear methods and the optimal algorithmic parameters. Our methods show that simple constrained matrix decomposition can produce to low-dimensional information in functionally interpretable and transferrable space, and can be widely useful for analyzing large-scale transcriptome data.

## Introduction

Recent developments in RNA-sequencing technology have enabled the collection of large-scale transcriptome data at high speed. For example, single-cell RNA-sequencing (scRNA-seq) data of many biological systems have been accumulating rapidly, and provide opportunities to gain insights into complex biological processes at both the systems level and the single-cell resolution. Together with the advances in experimental techniques, the recent development of computational methods, including those for dimensionality reduction, allow the visualization and analyses of high-dimensional transcriptome data in low-dimensional space. For example, t-distributed stochastic neighbor embedding (tSNE) and Uniform Manifold Approximation and Projection (UMAP) have been instrumental to tackling challenges in transcriptome data visualization and are widely used in biomedical research (1–4). However, dimensionality reduction methods usually do not provide low-dimensional space that is directly interpretable in terms of biological functions: while these approaches cluster related samples, the positioning of samples along the derived dimension does not correspond to the degree of any biological process even if a predefined gene set with similar functions is chosen before the reduction. In addition, the contribution or significance of individual genes regarding the derived dimension cannot be accessed directly with these methods. The lack of interpretability of the dimensions makes it challenging to visualize and analyze the progression of the samples (cells) in known biologically functional space.

Existing methods for functional quantification, such as z-score and Gene Set Variation Analysis (GSVA) (5), are useful for obtaining ‘functional scores’ with the expression levels of multiple genes involved in the same biological process. However, these methods do not have transferability in that the scoring systems obtained with one dataset cannot be used to analyze other datasets directly. This limits the utility of these methods in predicting the progress of new data points, and in studying the relationships between functional spaces in different experimental settings.

One example of cellular processes that contains crucial quantitative information is epithelial-mesenchymal transition (EMT). While extreme changes of cell fate and morphology occur in the classical form of EMT, recent studies with cancer and fibrosis showed that partial EMT involving intermediate states are prevalent, and it may be responsible for pathogenesis (6). To quantify the degree of EMT in EMT-induced cell lines and tumor samples, several previous studies analyzed transcriptomic data and their projections onto epithelial (E) and mesenchymal (M) dimensions (7–9). Recently, scRNA-seq analysis has shown that the progression of EMT is highly dependent on inducing signals and cell types (10). However, it remains challenging to analyze rapidly accumulating transcriptome information on EMT for obtaining biological insights across multiple conditions. Improvement of methods for reducing dimensions of expression data in a functionally meaningful manner is necessary.

In this study, we used gene set filtered variants of both non-negative principal component analysis (gsPCA) and non-negative matrix factorization (gsNMF) to analyze progression of EMT in single cells at multiple timepoints. We show that these methods describe large-scale transcriptome data of multiple EMT stages in low-dimensional and functionally interpretable space. Taking advantage of the methods’ transferability, we constructed dimensionality reduction models that can predict the stages of EMT with data from timepoints that were not used for model construction. We show that these linear methods can be used to compare functional spaces across multiple experimental conditions. Furthermore, we demonstrate the utilities of our approach in visualizing drug responses in heterogeneous single cell data. With a validation scheme for rigorous testing, we provide recommendations for the choice of the methods and the parametric settings. Overall, our work provides a new toolbox for analyzing large-scale transcriptome data with efficient visualization and functional quantification.

## Results

### Overview of method and performance evaluation

The overall goal of our method is to find low-dimensional space of transcriptome data that has both biologically meaningful directionality and the ability to represent data points not used in the procedure to derive the space. This requires one or more preselected functional gene sets, which are readily available in publicly available databases such as Molecular Signature Database (11), and can be defined manually (Figure 1). We propose two linear approaches of matrix decomposition: gsPCA and gsNMF (see Methods for details). Briefly, gsPCA finds the optimal component (projection) by maximizing the variance of the projected data points under the constraint that each functional gene has a nonnegative loading value. For gsNMF, the gene-set-filtered transcriptome matrix is approximated by the product of two nonnegative matrices, one of which represents the nonnegative weights of the functional genes (the procedure for obtaining the number of components is described in Supplemental Methods). Following gsNMF, the leading component is selected for subsequent analyses (see Methods). With either gsPCA or gsNMF, transcriptome data can be projected onto an axis whose direction unambiguously represents expression of the gene set and can be interrogated to reveal the contribution of individual genes in the set to scores along the axis.

**Figure 1.**
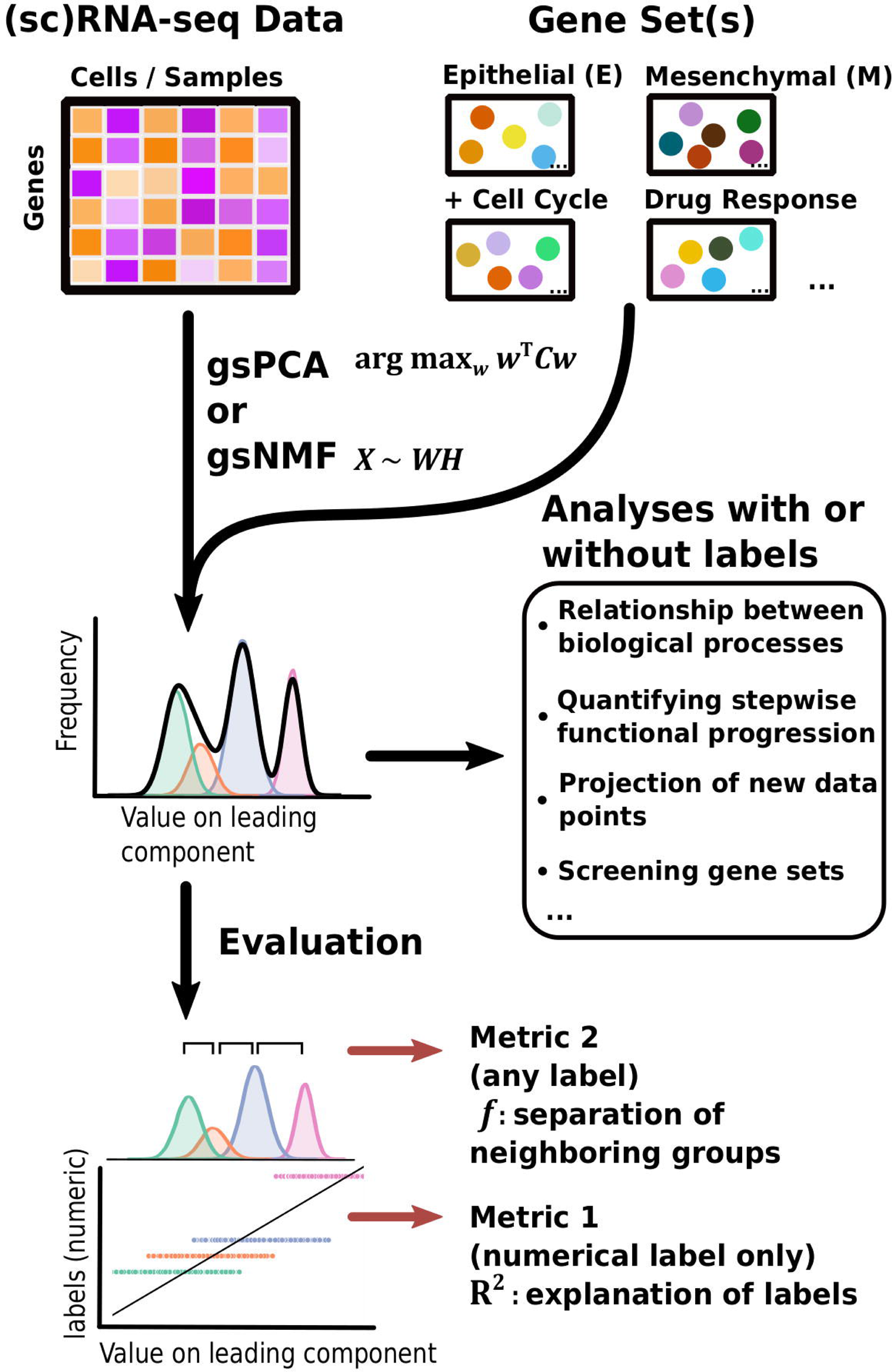
Schematic of the gsNMF/gsPCA analysis process. A diagram of the analysis process used in this study beginning with input data in the form of sequencing data and gene sets. gsNMF/gsPCA is applied to this data to generate a functional scoring or space in the form of component scores. These scores can be used in two ways. First, without further data labels, scores can be used to look at relationships between or across biological processes beginning with low-dimensional visualization to identify trends and putative groups, but also using quantitative approaches such as looking at the correlation between different functional scores. In addition, the transferable nature of these models means that they can be used to infer the position of new data points and the weighing of individual gene contributions allows for their importance to be assessed. Screening can be done both between gene sets and withing gene sets. Secondly, when data labels are present, different metrics can be used to assess the performance of a functional score in terms of capturing variance: the common language effect size or *f*-probability can be used to evaluate how well the functional score separates two distinct populations while the variance explained or R^2^ can evaluate how much of the variation of a linear, numeric variable, such as time, the functional score can explain across the data.

To test the performance of gsPCA and gsNMF in capturing biological progression through functional space, we first used time-course datasets containing single cells treated with EMT-inducing signals for various periods of time (10). In addition to the biological importance of the stepwise progression in EMT (6, 12), the time labels in the datasets allow us to evaluate the performance of the functional projection. Specifically, we used two metrics for the evaluation: the coefficient of determination (R^2^) for quantifying how well the projected values explain the time labels, and the common language effect size (*f*) for measuring the separation between two neighboring subsets of data with two labels (13) (See Methods). The usage of R^2^ is only possible when the labels are numerical, and *f* can be used with any type of label (Figure 1). Note that our goal is not clustering the data points. Instead, we aim to represent the progression along biologically meaningful axes. In addition, neither gsPCA nor gsNMF requires data labels for analysis. The two metrics are only used for evaluation. In later sections, we will show analyses with additional data sets in which labels are categorical and the biological processes are non-EMT.

### gsPCA and gsNMF capture cell state progression in low dimensional functional space

To show the performance of the proposed methods, we first used two signature gene sets whose high expressions represent the epithelial (E) and mesenchymal (M) states respectively (7, 9, 14). With the E and M gene sets, we first performed gsPCA and gsNMF on time-course single-cell transcriptomes of TGF-β-treated A549 cells using two components per model for each gene set (10). The two gene sets contain 176 and 116 genes, respectively, in the A549 data set. We then projected the single-cell data from the first five time points, which represent continuous EMT progression, onto the leading dimension for each gene set. This produced two-dimensional plots with dimensions that can be viewed as the progression of cell states in the epithelial and the mesenchymal spectrums (Figures 2A and 2B). We then compared the performance to two widely used approaches: z-score and GSVA (Figure 2C and 2D). We found that gsPCA and gsNMF both better explained the overall variance of time across the first five time points of EMT progression (Adjusted R^2^ = 0.46 and 0.48, respectively) than z-score (Adjusted R^2^ = 0.31) and GSVA (Adjusted R^2^ = 0.08). Likewise, when considering neighboring time points, we found that E-scores tended to decrease and M-scores tended to increase with time of TGF-β treatment (Figure 2E), with both scores significantly separating all neighboring time points for gsPCA and gsNMF and yielding higher *f* probabilities than other methods in all but one case (E-scores at 3 days vs 7 days, Figure 2F). This suggests that gsPCA and gsNMF not only serve as visualization methods of functional space with defined gene sets, but also describe heterogeneous cell populations containing transitional information in a rigorous fashion. Between the two methods, we found the gsNMF performed better with regard to both overall variance (Adjusted R^2^ 0.48 vs 0.46) and separating time points (Figure 2F) than gsPCA. However, gsNMF requires selecting the leading dimension based directly on correlation with time of EMT progression, suggesting that gsPCA may be more reliable in a purely unsupervised setting (see Methods).

**Figure 2.**
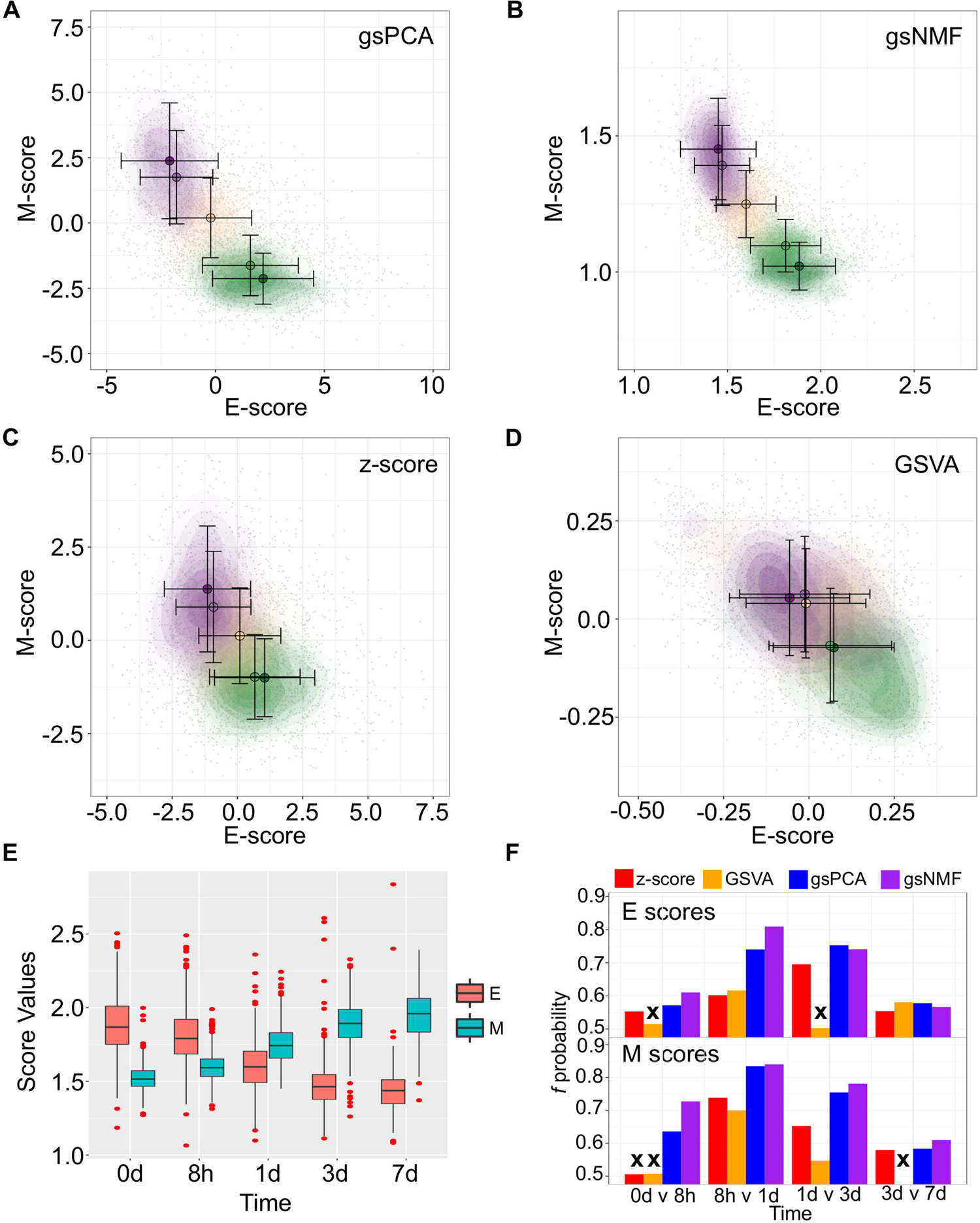
Visualization of EMT progression in TGF-β induced A549 cells by multiple scoring methods. (**A-D**) Contour plots of gene set scores of E (x-axis) and M (y-axis) genes from four different scoring methods, z-score (A), GSVA (B), gsPCA (C), and gsNMF (D). Color indicates the time of TGF-β induction from 0 days (dark green) to 7 days (dark purple). Circles indicate the mean E- and M-score of samples from each time point and the associated error bars show the standard deviation. (**E**) A box-plot showing the distribution of E (red) and M (blue) scores across all five time points of TGF-β induction from the gsNMF model. Whiskers indicate the 1.5 inter-quartile range of each distribution while the red points indicate outliers beyond this range. (**F**) Bar chart of the *f* probability values for E (top) and M (bottom) scores between all consecutive pairs of time points. Color indicates the method used to produce the score: red is z-score, orange is GSVA, blue is gsPCA, and purple is gsNMF. Bars marked by an ‘**x**’ indicates that the score did not significantly separate the samples from those time points (Mann-Whitney U-test, *p* < 0.05).

In the next few sections, we show various utilities of these linear methods based on their transferability and high-performance features. Because gsNMF gives the best performance with the A549 EMT data set, our discussion will focus on results obtained with gsNMF. The results using gsPCA, which had similar performance in all cases, are included in supplementary materials.

### Prediction of cell states with data from new conditions

The transferability of gsPCA and gsNMF methods allows the projection of new high-dimensional data points onto previously derived functional dimensions. Similarly, these methods can be used to derive functional dimensions with partial information of the biological process in terms of its stages. To show the predictive power of gsNMF, we removed samples from the 0-day, 1-day and 7-day (including revertant) time points in the A549 EMT data (i.e. the start, middle, and the end of the continuous portion of TGF-β induction) and then performed the dimensionality reduction. We found that the low-dimensional functional space was robust with respect to the removal, regardless of whether the missing time point is in the middle of the progress or at the extremes (Figure 3, A through D), such that when we projected the removed data points onto the space derived from a partial dataset, their positions were highly correlated with their positions when they were included in the data set (Pearson correlation coefficient (PCC > 0.95). However, while the inferred 1-day samples were similarly separable from samples in 8-hr (*f* = 0.81 for E, 0.84 for M) and 3-day (*f* = 0.75 for E, 0.79 for M) time points, we observed reduced separability between the both inferred 0-day vs. 8-hr (*f* = 0.52 for E, 0.63 for M) and 3-day vs inferred 7-day (*f* = 0.53 for E, 0.55 for M) time points, with E-scores not significantly separating the first and the last time points (Mann-Whitney U-test, p = 0.13 and 0.16, respectively). We also applied the same inference procedure to samples which were exposed to a transient EMT-inducing signal and allowed to revert. However, because the 8-hr and 24-hr reversion samples largely overlap with 7-day (hence their removal for 7-day inference, Figure 3E), we focused on inferring 3-day reversion samples after performing dimensionality reduction on the data set without any reversion samples. We found that 3-day reversion samples were positioned in the middle of the EMT spectrum, consistent with when they were included in functional space construction (PCC = 0.99 for E and 0.98 for M, Figure 3F). Additionally, the inferred 3-day reversion samples were similarly separable from the 7-day samples (*f* = 0.91 for E, 0.91 for M) as when they were when included in functional space construction (*f* = 0.90 for M, 0.91 for M). We obtained similar results using gsPCA when inferring the position of samples from missing time points (Figure S1), but neither E-nor M-scores significantly separated the end points (0-day vs 8-hr and 3-day vs 7day). These results suggest that gsPCA and gsNMF can predict cell states of new data without retraining the model, and that these methods can be used to predict new cell states that have not been observed directly, though it may be difficult to separate these samples when they are positioned the edge of the spectrum and/or when the new samples are closely related to existing samples.

**Figure 3.**
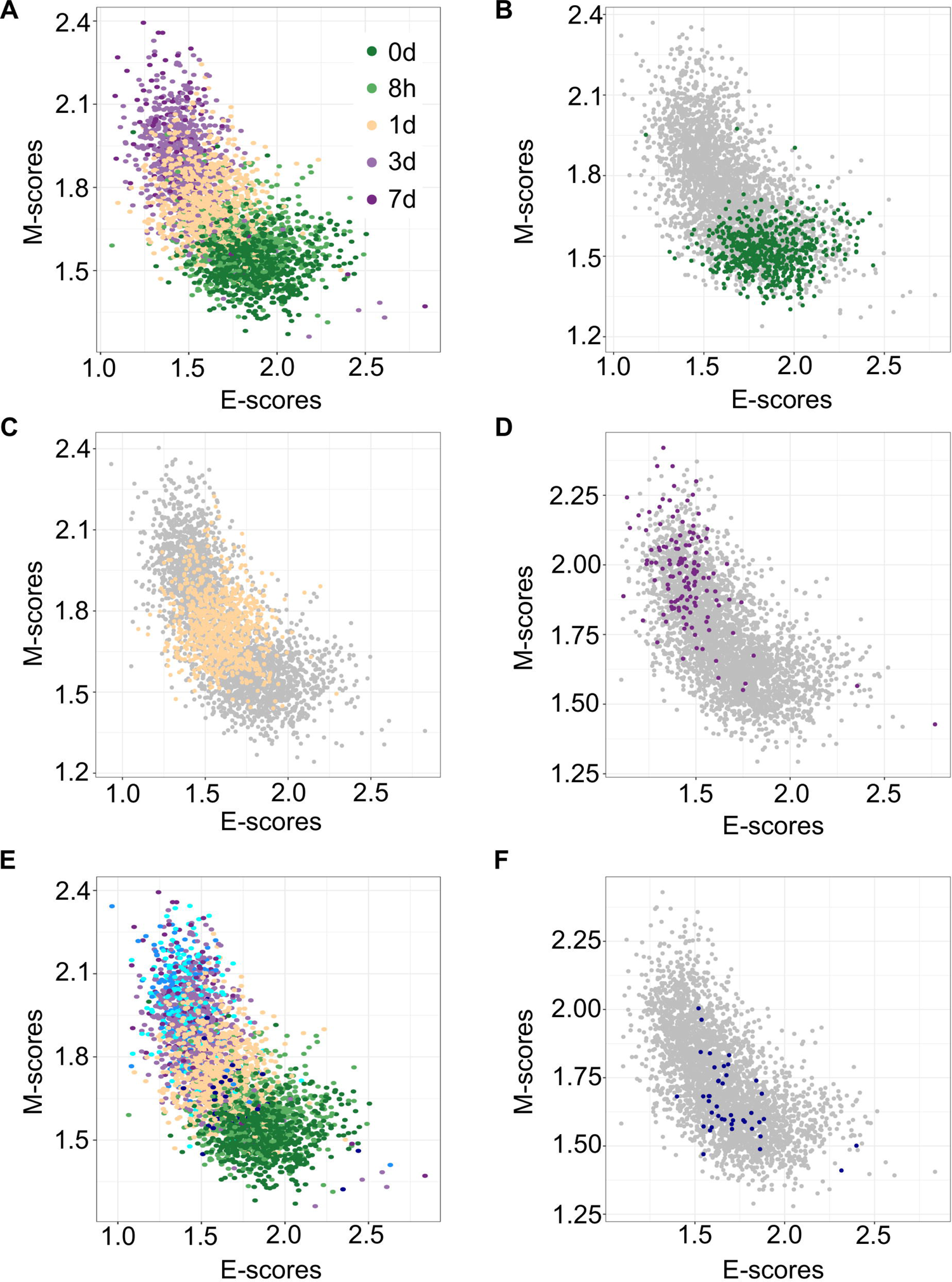
Predicting A549 sample from specific time points using gsNMF. (**A**) Scatter plot of E (x-axis) and M (y-axis) scores for all TGF-β induction samples using gsNMF. Samples from different time points are indicated by color going from 0 days (dark green) to 7 days (dark purple). (**B-D**) Scatter plot of 0-day (green, B), 1-day (yellow, C), and 7-day samples (purple, D) inferred using a gsNMF model built with all other time points (gray). (**E)** A scatter plot of TGF-β induction samples with TGF-β reversion samples (i.e. 7 days induction followed by removal from TGF-β). Induction samples are labeled as in (A), while reversion samples are colored blue, with darker shade indicating longer time since removal. (**F**) Scatter plot of 3-day reversion samples (dark blue) inferred using a gsNMF model built with all non-reversion time points (gray).

### Using functional space across cell lines

The transferability of gsNMF can be extended to data from different cell lines. We performed gsNMF on single-cell transcriptomes of TGF-β-treated DU145 from Cook and Vanderhyden (10) using the same procedure as A549 and obtained a moderate explanation of variance in time of EMT progression using the E and M dimensions (Adjusted R^2^ = 0.31). We then inferred the position of the five continuous time points in the A549 data set using the DU145 model and vice versa (Figure 4 A through D). Transferred models (DU145 on A549 and A549 on DU145) were able to separate the individual time points, but overall performance decreased as they can explain only part of the variance seen in the original models (Adjusted R^2^ = 0.30 for DU145 on A549 and 0.25 for A549 on DU145). Therefore, it was expected that the individual sample scores would be positively correlated between models along both the E (Figure 4, E and F) and M dimensions (Figure 4, G and H). However, while the correlations between all pairs of scores were significant (minimum *p* = 2.7e-73), the correlation between E-scores was weaker overall and worse for models of DU145 (PCC = 0.31) than models of A549 (PCC = 0.46). Comparably, the M-scores for both models of A549 (PCC = 0.84) and models of DU145 (PCC = 0.84) were more highly correlated and consistent between models. However, none of the sample scores between A549 and DU145 models were as correlated as inferred sample scores from missing point and the complete A549 model (PCC > 0.95). This suggests a reduced transferability across cells lines compared to within cells lines. In addition, across the data sets we used, changes along the M dimension were more consistent than the E dimension. We observed similar results using gsPCA, including M-scores being more correlated (PCC, A549 = 0.92, DU145 = 0.94) than E-scores (PCC, A549 = 0.76, DU145 = 0.72) (Figure S2). This is consistent with the fact that the same inducing agent was used across all cell lines, and also implies that inducing EMT in different cell types may yield more consistent changes in M genes compared to E genes.

**Figure 4.**
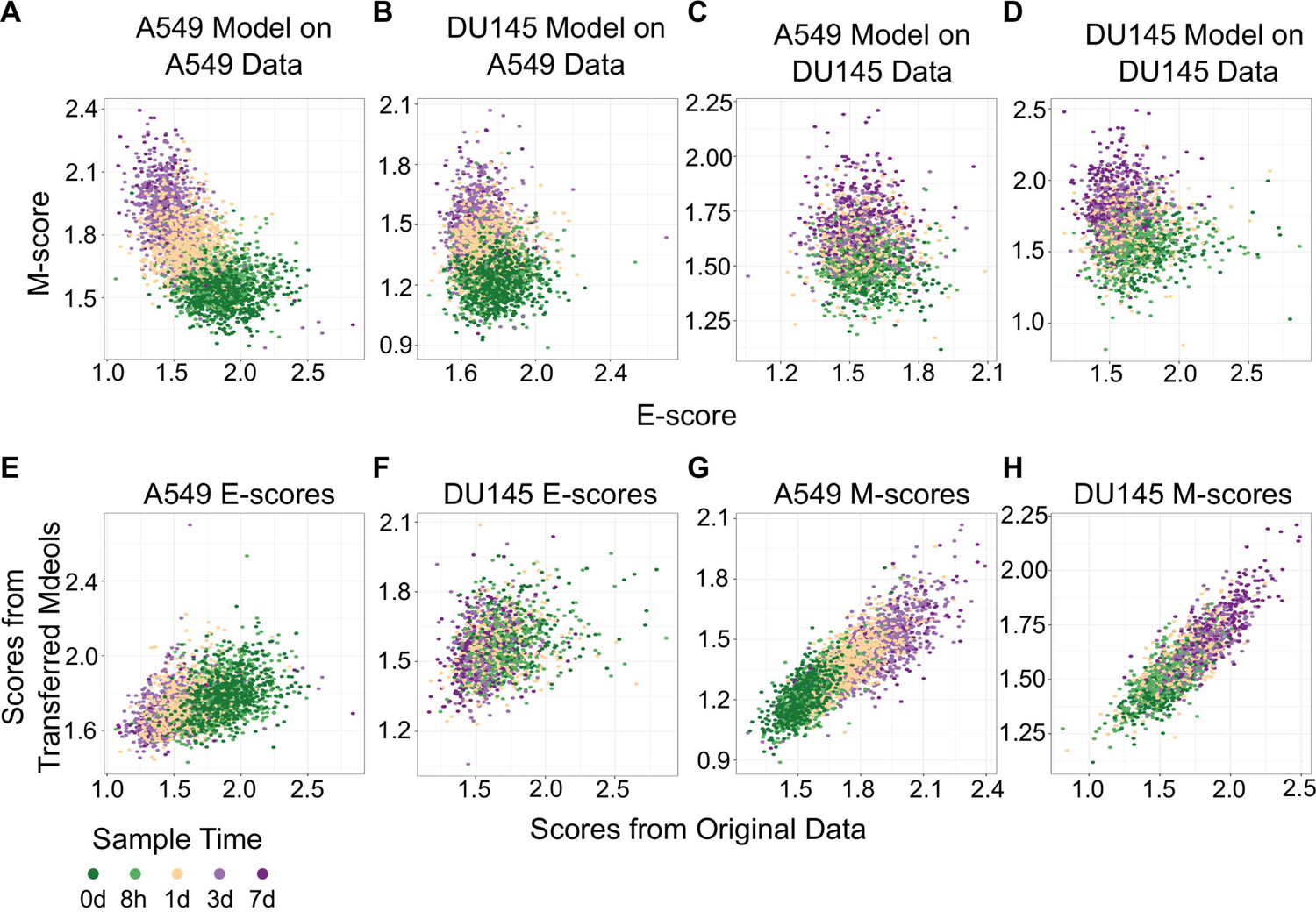
Transferring gsNMF models between A549 and DU145 TGF-β induced samples. (**A-D**) Scatter plot of E (x-axis) and M (y-axis) scores for different combination of data and gsNMF model: (A) A549 model on A549 data, (B) DU145 model on A549 data, (C) A549 model on DU145 data, and (D) DU145 model on DU145 data. Samples from different time points are indicated by color going from 0 days (dark green) to 7 days (dark purple). (**E-F**) Comparison of E-scores of samples from A549 (E) and DU145 (F) data. The x-axis is the E-score from using the model from the same data set (A549 on A549, DU145 by DU145), while the y-axis is the E-score from the opposite model (DU145 on A549, A549 on DU145). Samples from different time points are indicated by color going from 0 days (dark green) to 7 days (dark purple). (**G-H**) Comparison of M-scores of samples from A549 (G) and DU145 (H) data. The x-axis is the M-score from using the model from the same data set (A549 on A549, DU145 by DU145), while the y-axis is the M-score from the opposite model (DU145 on A549, A549 on DU145). Samples from different time points are indicated by color going from 0 days (dark green) to 7 days (dark purple).

### Using functional space across experimental conditions

In addition to predicting the locations in the functional space across cell lines, gsPCA and gsNMF can be used across both experimental conditions and cell types. To test the cross-condition transferability, we first used our low-dimensional functional EMT space for A549 and DU145 cells to analyze tumor transcriptomes measured with bulk RNA-seq (The Cancer Genome Atlas, TCGA). To perform the most comparable transfer, we used lung adenocarcinoma (LUAD) and prostate adenocarcinoma (PRAD) data, which correspond to A549 and DU145 in terms of tissue type. We considered transfers between both similar (projecting LUAD data by a A549-trained model, and PRAD by a DU145-trained model) and dissimilar (LUAD by DU145 and PRAD by A549) cell types. We found that the low-dimensional functional space obtained with in-vitro data captured tumor sample heterogeneity in the EMT spectrum when compared to our previous GSVA analysis of the same data (Figure 5). Overall, the original E- and M-scores were significantly correlated with the A549 models in all cases (smallest *p* = 2.8e-34). Models from both cell lines showed similar correlation with the original GSVA scores for both LUAD E-scores and LUAD M-scores as well as PRAD M-scores, but the DU145 model was better correlated with PRAD E-scores than A549 (Table 1) Furthermore, we once again observed that M-models built on A549 and DU145 data were more similar to each other that E-models (Table 1), consistent with our comparisons of A549 and DU145 directly. We obtained similar results with gsPCA (Figure S3), which showed greater overall correlation with GSVA scores, but like gsNMF showed reduced correlation for the A549 model of PRAD E-scores as well as stronger correlation between M-scores and E-scores (Table S1). Overall, this suggests that the robustness of derived functional spaces is broadly transferrable across data and cell types.

**Figure 5.**
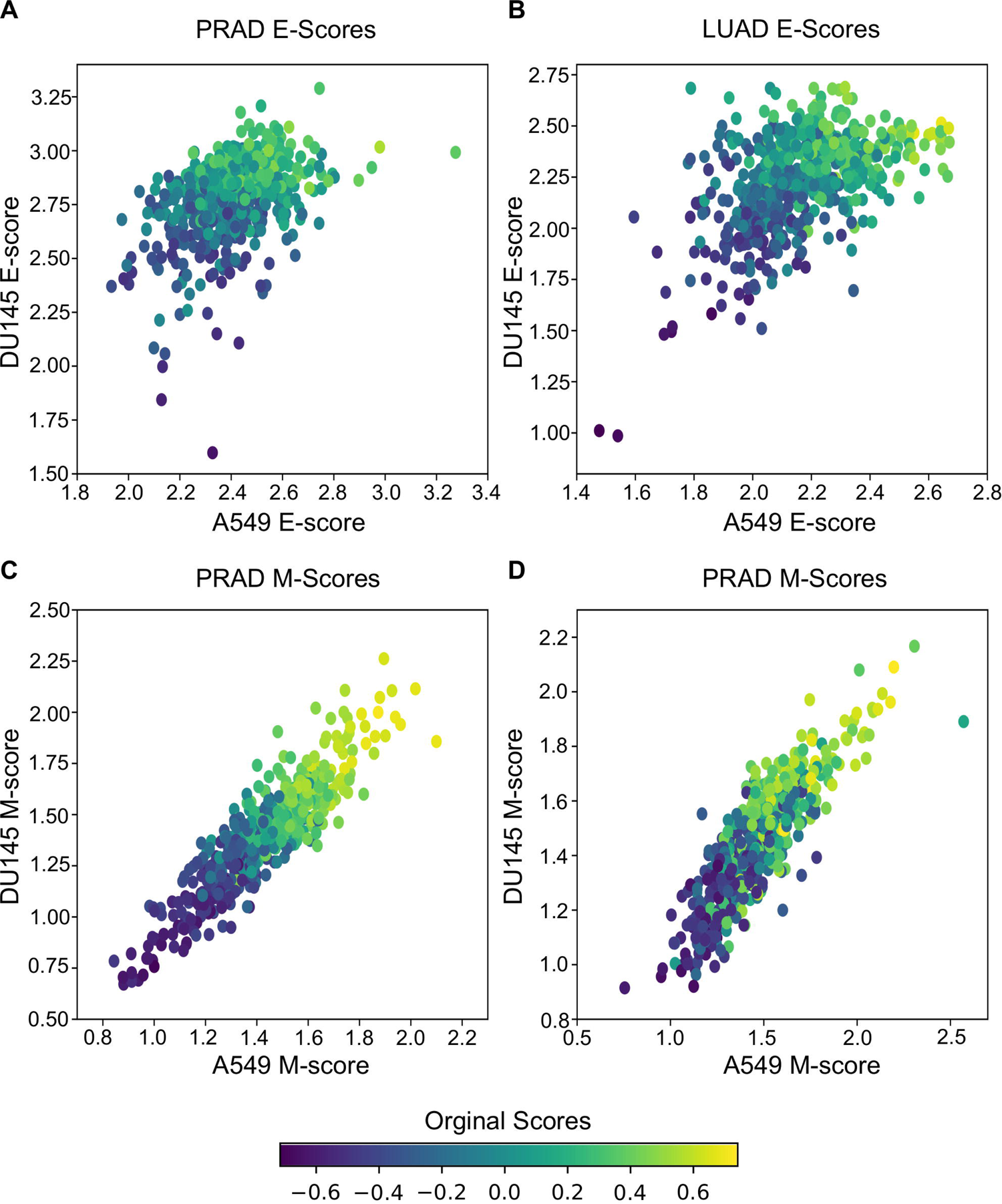
Transferring gsNMF models to TCGA data. (**A-B**) Scatter plots of E-scores for PRAD (A) and LUAD (B) from transferring gsNMF models built on A549 (x-axis) and DU145 (y-axis) data. The color of individual points indicates the original GSVA based E-score of the TCGA data set. (**C-D**) Scatter plots of M-scores for PRAD (C) and LUAD (D) from transferring gsNMF models built on A549 (x-axis) and DU145 (y-axis) data. The color of individual points indicates the original GSVA based M-score of the TCGA data set.

**Table 1.**
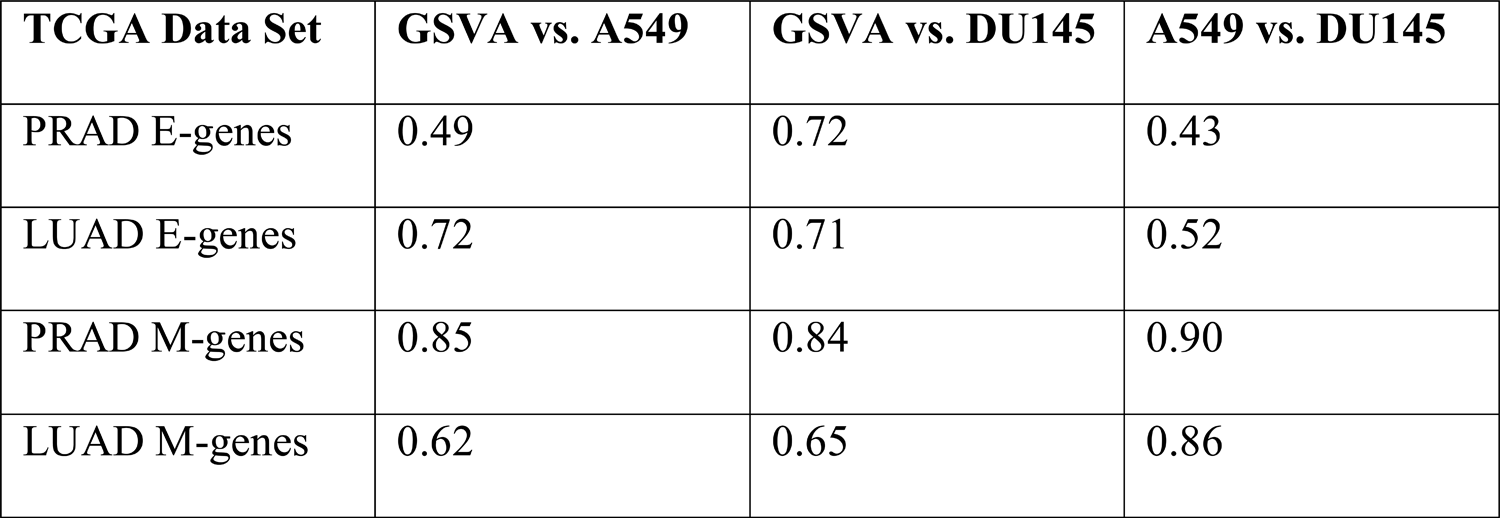
Pearson correlation coefficients of E and M scores between GSVA, A549 and DU145 Models of TCGA data

### Using functional space across spatial and temporal progression

We next examined if gsNMF can produce transferrable models that reveal both spatial and temporal progression of EMT. Using single-cell RNA-seq, McFaline-Figueroa et al. previously found that epithelial cells exhibit an E-to-M spectrum from the inner position of a colony to the outer position (15). This dataset that contains binarized identities (inner and outer) obtained with macro-dissection (defined as spatial EMT data) from two experiments, one in which cells were allowed to migrate without external β). Since there are only two populations in this data set, the leading dimension for E- and M-scores was chosen to maximize the separation based on the *f* probability. Overall, three analyses were performed for each data set: spatial data with its own gsNMF model, spatial data with the model from the other spatial data set (TGF-β on Mock and Mock on TGF-β), and spatial data with A549 time series model (Figure 6). As with our previous results, the best separation of inner and outer data points was observed when Mock (*f* = 0.61 for E, 0.73 for M) and TGF-β (*f* = 0.77 for E, 0.82 for M) data sets had their own model applied to them.

**Figure 6.**
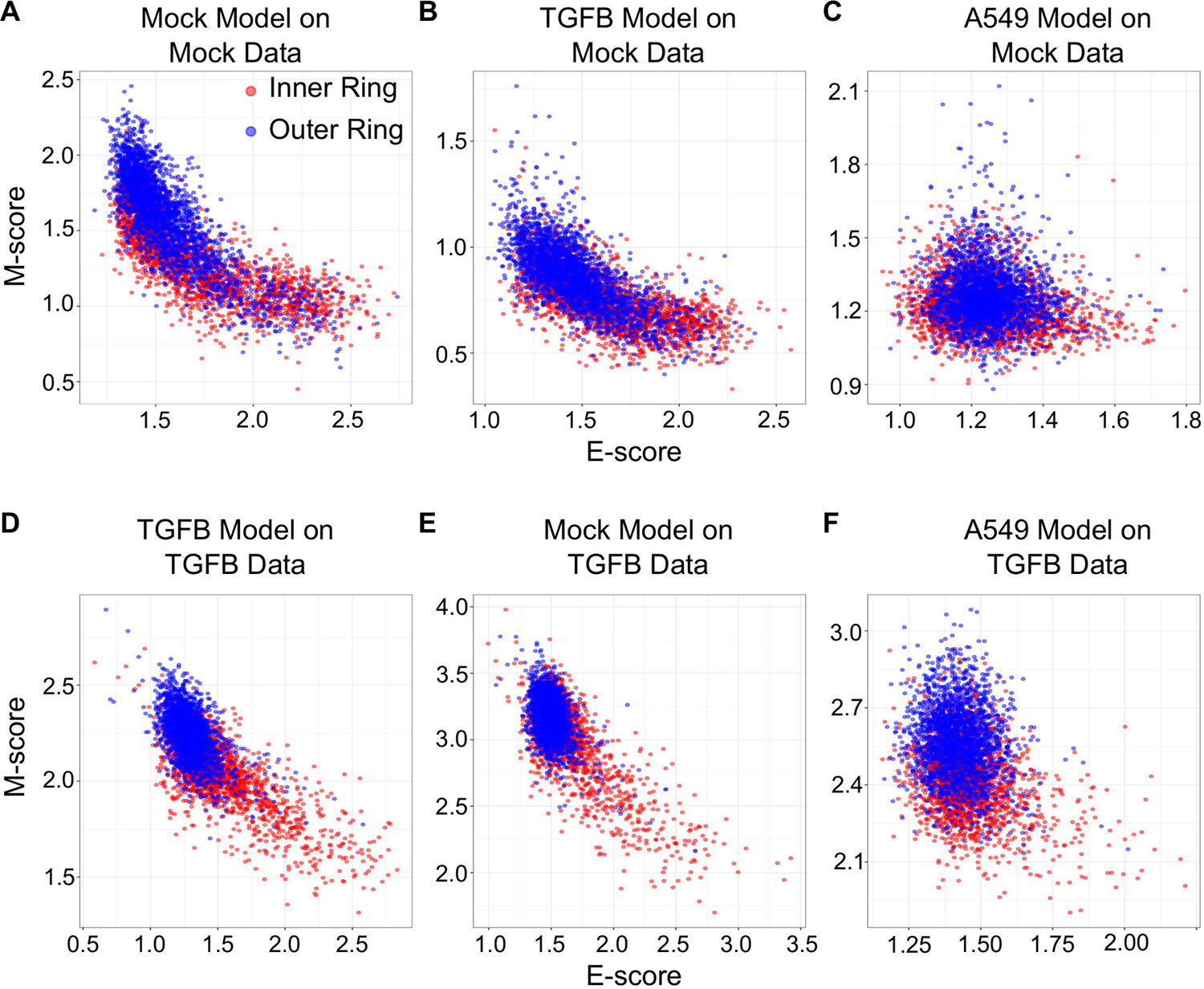
Transferring gsNMF models between temporal and spatial data sets. (**A-C**) Scatter plots of E (x-axis) and M (y-axis) scores for Mock spatial data from gsNMF models built on different data sets: Mock spatial data (A), TGF-β induced spatial data (B), and TGF-β induced A549 temporal data (C). The color of the sample indicates whether it originates from a cell in the inner-ring (non-motile, red) or the outer ring (motile, blue). (**D-F**) Scatter plots of E (x-axis) and M (y-axis) scores for TGF-β spatial data from gsNMF models built on different data sets: TGF-β induced spatial data (D), Mock spatial data (E), and TGF-β induced A549 temporal data (F). The color of the sample indicates whether it originates from a cell in the inner-ring (non-motile, red) or the outer ring (motile, blue).

However, for Mock data, the TGF-β model (*f* = 0.64 for E, 0.69 for M) outperformed the A549 model (*f* = 0.45 for E, 0.60 for M) on both dimensions and, in fact, the E dimension of the A549 model did not effectively separate inner and outer points in the Mock data (*p* = 0.99). In comparison, the Mock model better separated TGF-β inner and out points in the E direction (*f* = 0.68 for E, 0.63 for M), while the A549 model better separated them in the M direction (*f* = 0.59 for E, 0.76 for M). gsPCA models gave similar results, including the A549 model yielding better performance along the M-dimension (*f* = 0.76) for TGF-β data than Mock data (*f* = 0.68) (Figure S4).

The fact the A549 model better separated TGF-β spatial points along the M dimension than the Mock model, but did not outperform TGF-β on the Mock model suggests that there is conserved TGF-β induced M-gene expression regardless of context. To explore the basis of this similarity in M-scores, we compared the loading matrices between Mock, TGF-β, and A549 gsNMF models, which represent the weights of individual genes along the leading axis. We found little correlation between A549 and spatial E-gene loading along the lead dimensions (PCC = −0.02. *p* = 0.82 for Mock; PCC = 0.06, *p* = 0.57 for TGF-β), however, while there was also little correlation between A549 and spatial M-gene loading values for the Mock model (PCC = 0.02, *p* = 0.83) there was significantly positive correlation for the TGF-β model (PCC = 0.48, *p* = 8.8e-7). Additionally, we examined which genes were in the top 10^th^ percentile of loading values across models and found that the A549 and TGF-β models share six M-genes (FN1, LGALS1, SERPINE1, TAGLN, TPM2, VIM), compared to three E-genes (ELF3, PERP, SLPI).

Furthermore, another four E-genes (’AREG, KRT18, KRT8, NQO1) were in the top 10^th^ percentile of A549 E-gene loading, but the bottom 10^th^ percentile of TGF-β E-gene loadings. We observed similar results from gsPCA, finding significant correlation loading values only between A549 and TGF-βM-models (PCC = 0.61, *p* = 5.8e-11) with many of the same genes in the top 10^th^ percentiles of both models (FN1, TPM2, VIM, TAGLN, GLIPR1, LGALS1). Together, these results suggest a coherence of the progression of EMT program in both the spatial and temporal context with regard to M-genes, while E-gene progression appears to be more sensitive to context, being only transferable between the two spatial data sets. In addition, they highlight the usefulness of the transferability of gsPCA and gsNMF outside of a time series context, where performance may need to be evaluated on discrete groups.

### Characterizing relationships among multiple functional spectrums

To test the capacity of gsNMF to infer functional spaces across a broader range of gene sets and data, we first returned to the A549 data set and examined the expression changes of multiple gene sets across EMT progression. Taking advantage of the high-efficiency of this method, we began with 5455 C2 curated gene sets from the Molecular Signature Database (see Methods) and applied a gsNMF model to A549 data for each. For simplicity, we used a two-component model, but we applied stricter convergence and selection criteria because of the diversity of gene set size and coverage by the data set (see Methods). Overall, 867 gene sets (15.9%) had a leading dimension whose magnitude of correlation (PCC) was > 0.5 (Table S2). As such, we expected that functional spaces constructed from highly correlated gene sets should show similar results to our original E vs. M functional space.

To construct unambiguous functional spaces, we initially focused on pairs of up/down regulated gene sets where the leading dimensions had a high magnitude of correlation (PCC), but opposite sign, in order to emulate our original E/M model of EMT progression for A549 (Figure 7A). For example, two pairs of gene sets, up regulation or down regulation in response to KRAS knockdown (SWEET_KRAS_TARGETS, Figure 7B) and up regulation or down regulation in low-malignancy ovarian cancer relative to control (WAMUNYOKOLI_OVARIAN_CANCER_LMP, Figure 7C), yielded functional spaces similar to E and M genes (Figure 7A) and captured a similar amount of variance explained among non-revertant cells (R^2^ = 0.48 and 0.49, respectively). Furthermore, the results suggest that EMT progression is correlated with expression of gene normally repressed by KRAS, a pro-proliferation signal, and anti-correlated with the expression of genes associated with tumorigenic, but non metastatic ovarian cancer, consistent with the idea of the E state of EMT being pro-proliferative and the M state being pro-migratory. However, not all pairs of gene sets provide strongly functional spaces: for example, the gene set down-regulated in metastatic vs. non-metastatic head and neck tumors (RICKMAN_METASTASIS_DN) produced a strong anti-correlated leading dimension (PCC = −0.63), but the leading dimension of the up-regulated variant has a far smaller magnitude of correlation (PCC = 0.34). However, combining the metastatic down-regulated gene set with another correlated gene set, genes silenced during angiogenesis (HELLEBREKERS_SILENCED_DURING_TUMOR_ANGIOGENESIS, PCC = 0.66), generated a functional space of EMT progression competitive with E and M genes (Figure 7D, R^2^ = 0.48). As such, functional space constructs need not be confined to reciprocal or connected gene sets, though the enrichment of EMT genes suggests that there is probably an underlying, common genetic basis explain the similarity of all these functional spaces. Nevertheless, the divergent origins of the gene sets in terms of the biological processes they represent demonstrates the breadth over which the functional significance of variation can be explored using this methodology.

**Figure 7.**
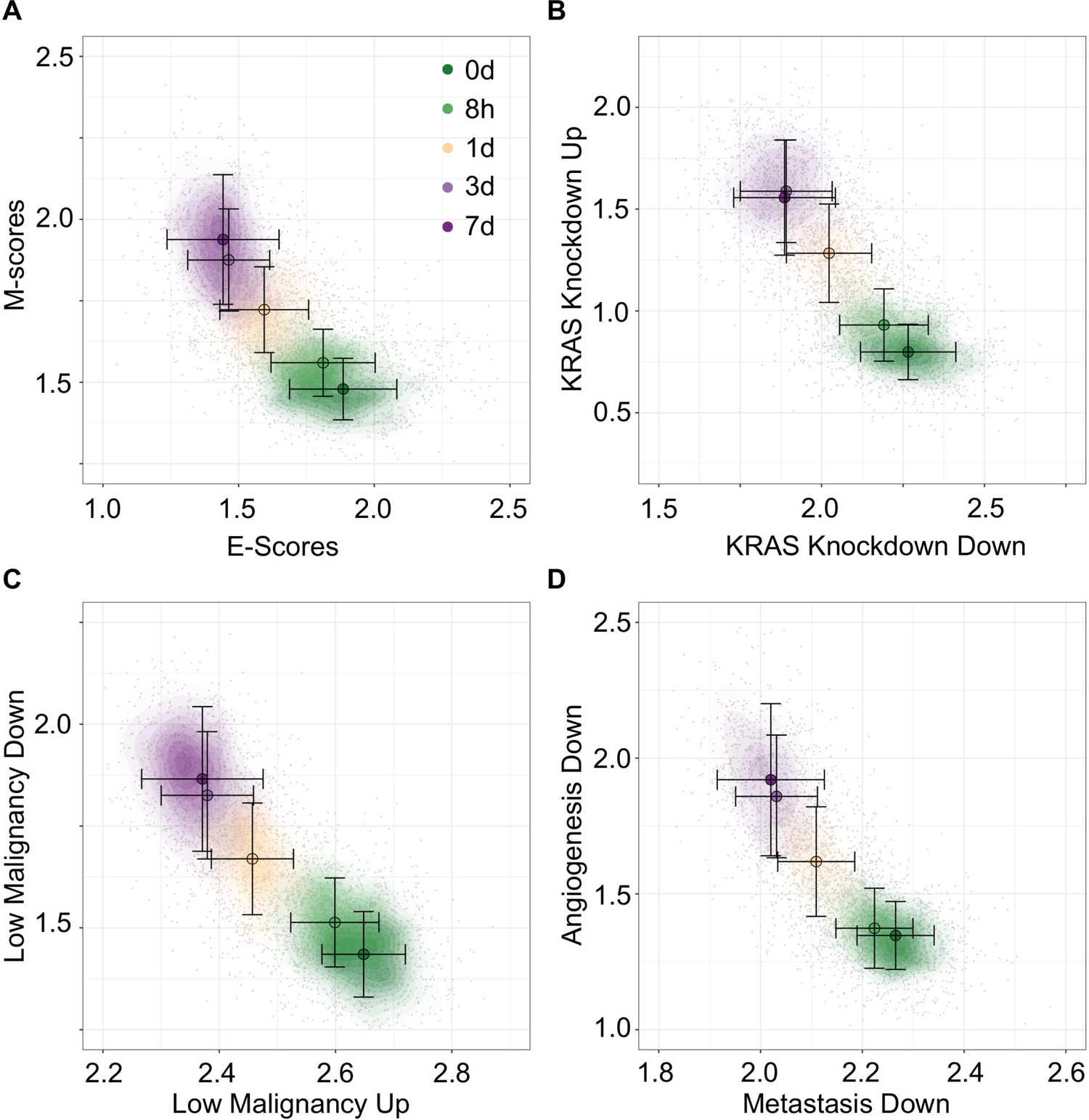
Visualization of EMT progression in TGF-β induced A549 cells by multiple gene sets. (**A-D**) Contour plots of A549 functional space generated using gsNMF with different gene sets: E vs M (A), KRAS knockdown up and down (B), non-malignant ovarian cancer up and down (C), and metastasis downregulation vs angiogenesis downregulation (D). Color indicates the time of TGF-β induction from 0 days (dark green) to 7 days (dark purple). Circles indicate the mean gene set score of samples from each time point and the associated error bars show the standard deviation.

To move beyond EMT associated data and gene sets, we next used gsNMF to analyze data from McFarland et al. (16) which is composed of 7245 cells with heterogeneous origins treated with trametinib for 3, 6, 12, 24,or 48 hours as well as an untreated control (0 hours). Because this data mixes sample from 24 cell lines from several different origin tissues and focuses specifically on the response to a cancer drug, we focused our exploration of functional spaces on 1022 gene sets derived from the C6 database from Molecular Signature Database as well as the drug resistant genes identified by Wang et al. (17) and their overlapping KEGG pathways and GO terms (see Methods). Overall, 57 gene sets (5.6%) had a leading dimension whose magnitude of correlation (PCC) was > 0.5 and relaxing this threshold to > 0.4 yielded only 200 (19.6%) of gene sets, suggesting that the explained temporal variance in this data set is lower than that obtained with A549 (Table S3). Nevertheless, using positive regulation of gene expression (GO:0010628) and negative regulation of gene expression (GO:0010629), we were able to a functional space of trametinib response with similar performance (R^2^ = 0.30, Figure 8A) our model of EMT progression in DU145 data (R^2^ = 0.31). Additionally, a number of oncogenic signatures which were positively correlated with trametinib response, though there were no up/down regulated pairs that with strong leading dimensions in both directions. Instead, we selected two oncogenic signatures, down regulation ins response to KRAS over-expression (KRAS.600_UP.V1_DN) and down regulation in response to LEF over-expression (LEF1_UP.V1), whose leading dimension were strongly correlated with trametinib response (PCC = 0.54). We then took the negatively correlated dimension corresponding up regulation gene sets (KRAS.600_UP.V1_UP and LEF_UP.V1_UP) which had similar magnitude of correlation along both dimensions (difference in PCC <= 0.005). This process gave functional spaces which improved variance explained over the previous gene regulation model (R^2^ = 0.35 and 0.36, respectively, Figure 8B and C). Together, these results suggest suppression of gene expression in general and of oncogenes in specific in response to trametinib treatment, consistent with the results in McFarland et al. which observed greater enrichment of KRAS responsive genes among down-regulated genes in later time points relative to earlier ones. As with A549 data, we were also able to combine distinct functional sets, response to drug (GO:0042493, PCC = 0.58) and positive regulation of cell cycle (GO:0045787, PCC = −0.49) to explain an comparable amount of variance in expression as the reciprocal onco-gene sets (R^2^ = 0.36, Figure 8D). As such, while the variance we can capture is dependent on the data set, our method is capable of producing functional spaces broadly capture variance in expression across diverse data and gene sets.

**Figure 8.**
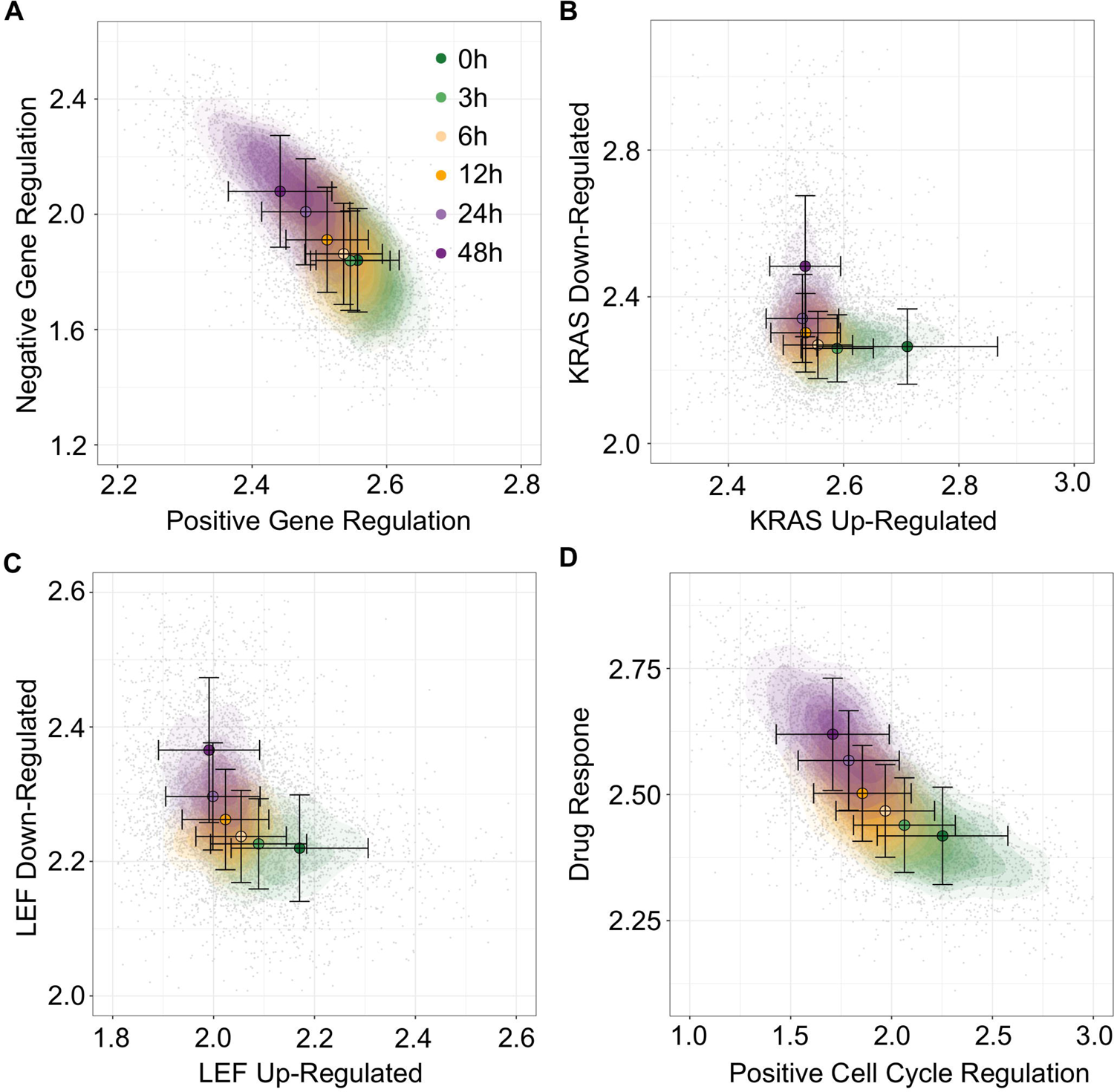
Visualization of trametinib treatment data by multiple gene sets. (**A-D**) Contour plots of trametinib treatment functional space generated using gsNMF with different gene sets: positive vs negative gene regulation (A), KRAS overexpression up and down regulation (B), LEF overexpression up and down regulation (C), and positive cell-cycle regulation vs drug response (D). Color indicates the time of trametinib treatment from 0 hours (dark green) to 48 hours (dark purple). Circles indicate the mean gene set score of samples from each time point and the associated error bars show the standard deviation.

## Discussion

Previous methods that aimed to address the challenges of visualizing single-cell data in functional space were primarily based on weighted sum of expression values or Kolmogorov–Smirnov test with full datasets (5, 18). These methods are useful to analyzing samples with functional gene sets, they do not provide transferability which is essential for predicting cell states with existing models and new data. We showed that constrained linear transformation enables good performance in depicting cell states with straightforward interpretation in functional space and satisfactory efficiency. While more sophisticated methods such as deep generative models have potentials to address similar problems, current methods primarily focus on the interpretability in terms of inter-sample distances in low dimensions rather than the dimensions themselves (19, 20), and we expect that the gsPCA and NMF methods are more efficient than models based on nonlinear connectivity.

Factorization approaches like PCA and NMF have previously been applied to the problem of gene expression, with NMF in particular having been used to deconvolute expression patterns scRNA-seq data sets (21, 22), but these approaches have primarily focused on the unsupervised clustering of samples and/or for *de-novo* module discoveries at relatively high dimensionality (n > 10). In contrast, our approach suggests there is a utility in applying these factorization approaches to interrogating the relationship between known gene modules and data with implicit structure and/or separable populations of samples, particularly when assessing a single biological process (EMT) across multiple contexts (e.g. cell line, time and space), such that the simplicity of low-dimension space (n = 2) can be leveraged for visualization and analysis.

In this work we have found that conserved EMT gene expression signatures can be used to describe stages of EMT in multiple cell lines (e.g. A549 and DU145), and these signatures not only capture the subpopulation heterogeneity resulting from differential times of treatment with EMT-inducing signals such as TGF-β, but also reflect the EMT program driven by spatial heterogeneity with cell populations (15). These results are consistent with the existence of conversed EMT program across cell lines (10), but do not contradict the idea of context specific expression as models trained and applied to the same data set always explained more variance in EMT progression. The coexistence of a common EMT signature and context specific expression is further supported by the observation that M-scores were more consistent and better separate data across different contexts of EMT than E-scores, and the related observation that M-gene loading values were correlated across spatial and temporal models, while E-genes were not. This suggests that M-gene induction by TGF-β is consistent across cellular contexts, while changes in E-gene expression are more variable, possibly due to greater sensitivity to cell line, environmental context, or other initial conditions effecting the cell prior to induction.

The transferability of models across EMT context indicates the synergy between spatial arrangement of cells and external signals (e.g. TGF-β) in determining the stages of EMT. In addition, we found that the functional dimensions obtained with TGF-β can serve as reasonable approximations for the positioning of tumor transcriptomes in the EMT spectrum. Similar to the EMT spectrum, many biological processes involve stepwise changes of gene expression programs. A possible mechanism underlying these non-binary programs is the feedback-driven formation of stable intermediate cell states (23, 24). With the rapid advances of the single-cell technology, transcriptome-wide gene expression data will become available for more biological systems. We expect that our functional projection methods can be widely useful for visualizing and analyzing these data. In particular, the transferability of the models can be a powerful feature for interrogating the relationships among different experimental conditions and cell types.

## Methods

### Gene expression data sources

scRNA-sequencing data and meta data for A549 and DU145 cell lines were obtained from Cook and Vanderhyden (10). In brief, we obtained pre-processed SeuratObject for A549 and DU145 TGF-β as .rds data files and extracted expression data for E, M, all genes using the ScaleData function from Seurat to regress out mitochondrial gene expression, total unique reads in a sample, cell cycle gene expression, and batch effects as well as scale each data set across genes (25). For McFaline-Figueroa et al. spatial data we obtained aggregated count data from GEO in the form of a pre-processed .cds file (GSE114687, McFaline-Figueroa et al., 2019). We then dropped genes expressed in less than 50 cells (∼1% of each data set) from Mock and TGFB1 and split samples into Mock and TGFB1 subset for subsequent steps. Because we planned to compare models from these data to those from A549 and DU145, we followed the preprocessing procedure from Cook and Vanderhyden: we normalized the Mock and TGFB1 data sets independently in Seurat using the NormalizeData function and then used ScaleData to regress out mitochondrial gene expression, total unique reads in a sample, and cell cycle gene expression as well as scale each data set across genes. Finally, we obtained Cell Ranger output for trametinib time course data from McFarland et al. (16) and processed it in R using Read10X. We used annotations from the original manuscript to eliminate low quality cells and then filtered genes expressed in less than 73 cells (∼1% of the data set). Pre-processing was done in Seurat as with using NormalizeData and ScaleData as previously described, except that we additionally regressed out the effect of each different cell lines used in the experiment, but did not regress out cell-cycle gene expression as the original manuscript suggested that cell cycle disruption may be induced by trametinib treatment.

TCGA bulk RNA-seq data was obtained from TCGAbiolinks (15, 26). Raw counts were transformed to log_2_TPM with a pseudo-count of 1 using gene models for the hg38 annotation of the human genome obtained from RefSeq (27).

### Dimension reduction approaches

We implemented non-negative PCA in R using the *nsprcomp* function (with the option nneg=TRUE) from the package of the same name (28). We used the standard convergence parameters for the algorithm as these produced consistent principle components across multiple runs and different number of components. This is to be expected as *nsprcomp* greedily maximizes the variation explained by each component in order. For gsNMF, we used the Scikit-learn implementation of NMF (29). To optimize convergence criteria, we performed a cross-validation analysis of A549 data and found that a two component model fit with a tolerance of 1e-6 and a max of 500 iterations gave the best results (see Supplemental Methods and Figure S5). We also tested ten random seeds of the two component A549 model on the full data to confirm that consistent results were given (average PCC of dimensions > 0.99). We tested ten random seeds against the other data sets to tune the convergence parameters, raising maximum iterations to 2500 and tolerance to 1e-9 if the initial parameters did not yield consistent results (i.e. average PCC of dimensions > 0.99). Further background information about non-negative PCA and NMF can be found in the Supplementary Methods. GSVA and z-score methods were implemented using the GSVA package in R (5).

Unlike GSVA and z-scores methods, which produce a single score per gene set, gsPCA and gsNMF both produce multiple scores in the form of principle components (gsPCA) or the columns of the transform matrix (gsNMF). Therefore, we need to choose a ‘leading dimension’ to represent each gene set in functional space. For gsPCA, we use the first principle component as this represent the direction of greatest variance for gene expression in that gene set. For gsNMF, we used the magnitude of correlation between the columns of the transform matrix and the time of each sample to select a leading dimension. For E and M gene sets on EMT data, we also required the sign of correlation to match the expected change in E and M genes during EMT (i.e. picking the greatest negative PCC for E and the greatest positive PCC for M). For our spatial EMT data, where there were only two populations, *f* probability was used instead (see below), but with the same constraint on the direction of E and M dimensions (i.e. higher M scores for outer samples and higher E scores for inner samples).

### Evaluating functional spaces

To evaluate a functional space, we used two metrics. First, if the data had an associated time variable, we created a model of time as a linear function of the two axes of the functional space (time ∼ X + Y) and calculated the coefficient of determination, which is the percent of overall variance in the dependent variable explained by the independent variable (Adjsuted R^2^). Second, to evaluate the ability of functional space to separate distinct populations, such as neighboring time points or location, we used the common language effect size (*f*), which is the probability that if a pair of samples are chosen at random from two populations, the score of the sample from the ‘first’ population is higher. The metric is advantageous because we can calculate it from the test statistic of Mann Whitney U-test, which also provide a measure of significance, and is related to the area under of the receiver operating curve (AUC-ROC), which is common used to asses classification algorithms. Additionally, since the *f* probability is reciprocal, the choice of the ‘first’ population is arbitrary, so for EMT we can chose to calculate the *f* probability such that the ‘first’ population is the more progressed for M and less progressed for E. Therefore, a higher probability always indicates better correspondence with EMT progression for those data sets.

### Inference and model transfer

To infer the position of new data in functional space for gsPCA, we multiplied the new data directly by weight vectors (also known as rotation vectors) of the E- and M-scores. For gsNMF, we used the Scikit-learn ‘transform’ method which transforms the input data according to the fitted model. In both cases, we used the same leading dimension for inference as in the original model. For inferring missing data points, no further steps were required as the new data always had the same coverage of the E and M gene sets as the original. However, for transferring models across cell-line, TCGA, and spatial data, we first had to determine the common set of genes between the two data sets. Common genes were then used to filter the weight vectors for gsPCA and to refit the model to be transferred on the common subset of genes for gsNMF. Input data was then filtered by the same subset of data and inference was done as described previously. Transferred models were assessed against the new data set using the same approaches as the original models, but relationship between E/M-scores and gene rotation/loading values between models built on the original data and transferred models were assessed by PCC for both magnitude and direction.

### Multi-gene set evaluation

C2 gene sets were obtained from the Molecular Signature Database (version 7.1, http://www.gsea-msigdb.org/gsea/msigdb/index.jsp) (11, 30, 31). gsNMF was performed as described for EMT gene sets expect that we increased the iteration (2500) and convergence threshold (1e-9) of the NMF algorithm to ensure consistent results across the gene sets which varied widely in size (2 to 1581 genes present in the data set) and coverage by the A549 data set due to the sparsity of scRNA-seq data. To test the robustness of this approach, we looked at the correlation of PCC scores along the leading axis for each gene set across 10 random seeds and found they were highly similar (average PCC between seeds = 0.998).

We used the same iteration and convergence threshold for analysis of the C6 (Molecular Signature Database) and the GEAR drug resistance gene set (17) which were used to project the trametinib data. Gene sets, KEGG pathways and GO terms associated with the GEAR drug genes were obtained using KEGGREST package in R for KEGG pathways and http://geneontology.org/ for GO terms (32, 33).

## Data and Code availability

Code and data for generating the primarily results of this study can be found at https://github.com/panchyni/gsNMF

## Author contributions

Designed research: TH. Performed research: NP and TH. Analyzed data: NP, KW, and TH. Wrote manuscript: N.P. and T.H.

## Conflict of Interest

The authors declare no conflict of interest.

---

**Supplemental Figure S1.**
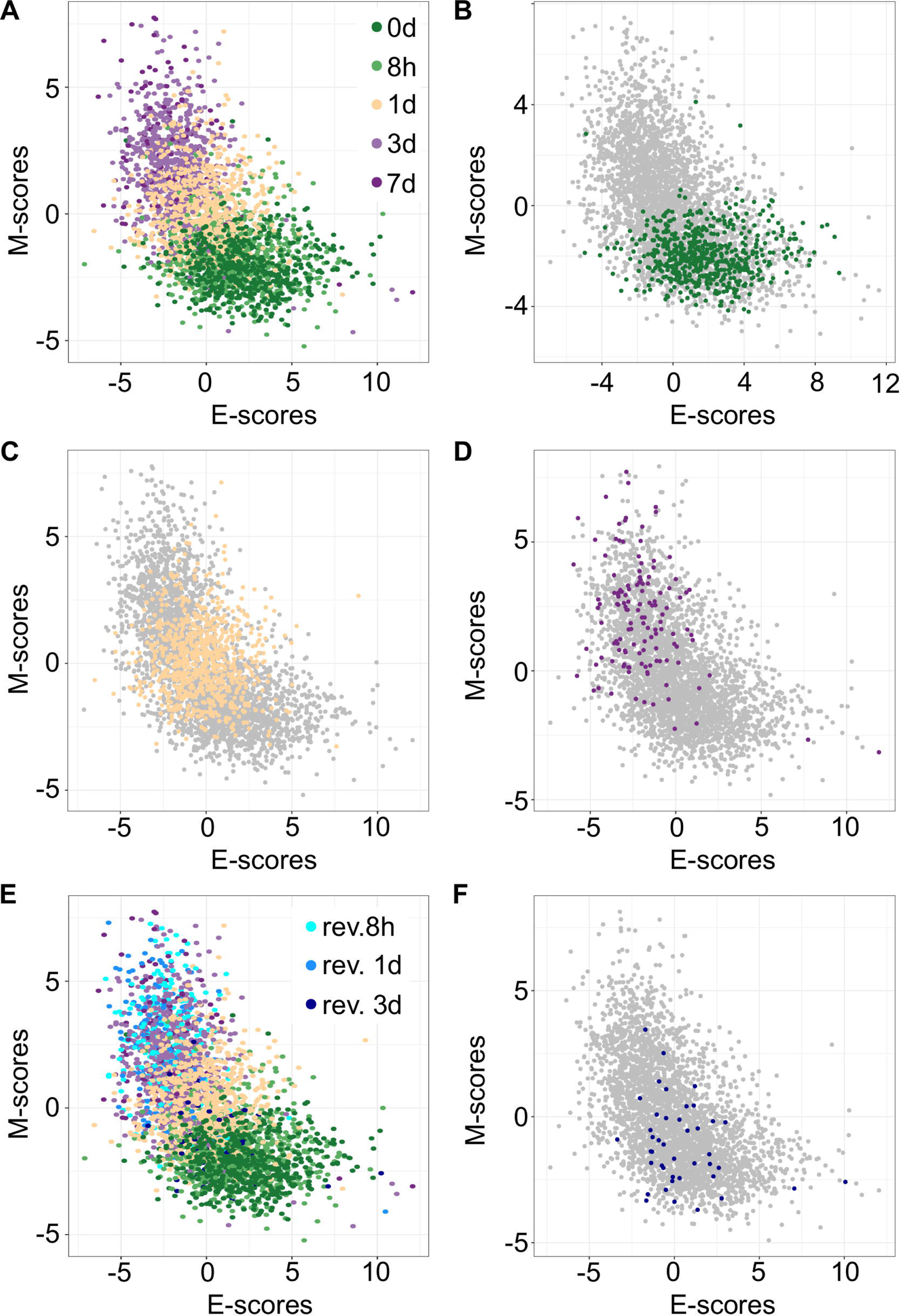
Predicting A549 sample from specific time points using gsPCA. (**A**) Scatter plot of E (x-axis) and M (y-axis) scores for all TGF-β induction samples using gsPCA. Samples from different time points are indicated by color going from 0 days (dark green) to 7 days (dark purple). (**B-D**) Scatter plot of 0-day (green, B), 1-day (yellow, C), and 7-day samples (purple, D) inferred using a gsPCA model built using all other time points (gray). (**E)** A scatter plot of TGF-β induction samples with TGF-β reversion samples (i.e. 7 days induction followed by removal from TGF-β). Induction samples are labeled as in (A), while reversion samples are colored blue, with darker shade indicating longer time since removal. (**F**) Scatter plot of 3-day reversion samples (dark blue) inferred using a gsPCA model built using all non-reversion time points (gray).

**Supplemental Figure S2.**
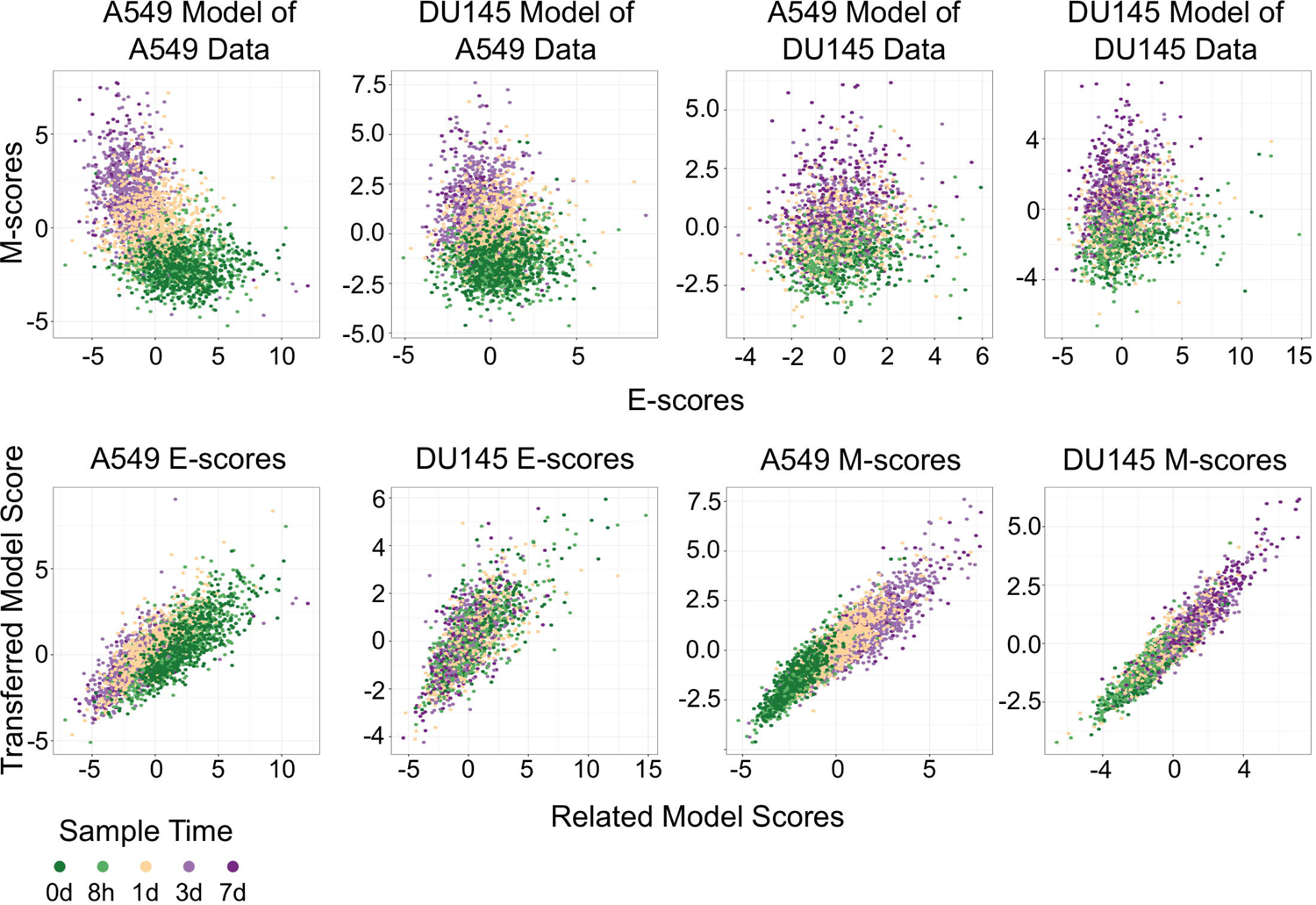
Transferring gsPCA models between A549 and DU145 TGF-β induced samples. (**A-D**) Scatter plot of E (x-axis) and M (y-axis) scores for different combination of data and gsPCA model: (A) A549 model on A549 data, (B) DU145 model on A549 data, (C) A549 model on DU145 data, and (D) DU145 model on DU145 data. Samples from different time points are indicated by color going from 0 days (dark green) to 7 days (dark purple). (**E-F**) Comparison of E-scores of samples from A549 (E) and DU145 (F) data. The x-axis is the E-score from using the model from the same data set (A549 on A549, DU145 by DU145), while the y-axis is the E-score from the opposite model (DU145 on A549, A549 on DU145). Samples from different time points are indicated by color going from 0 days (dark green) to 7 days (dark purple). (**G-H**) Comparison of M-scores of samples from A549 (G) and DU145 (H) data. The x-axis is the M-score from using the model from the same data set (A549 on A549, DU145 by DU145), while the y-axis is the M-score from the opposite model (DU145 on A549, A549 on DU145). Samples from different time points are indicated by color going from 0 days (dark green) to 7 days (dark purple).

**Supplemental Figure S3.**
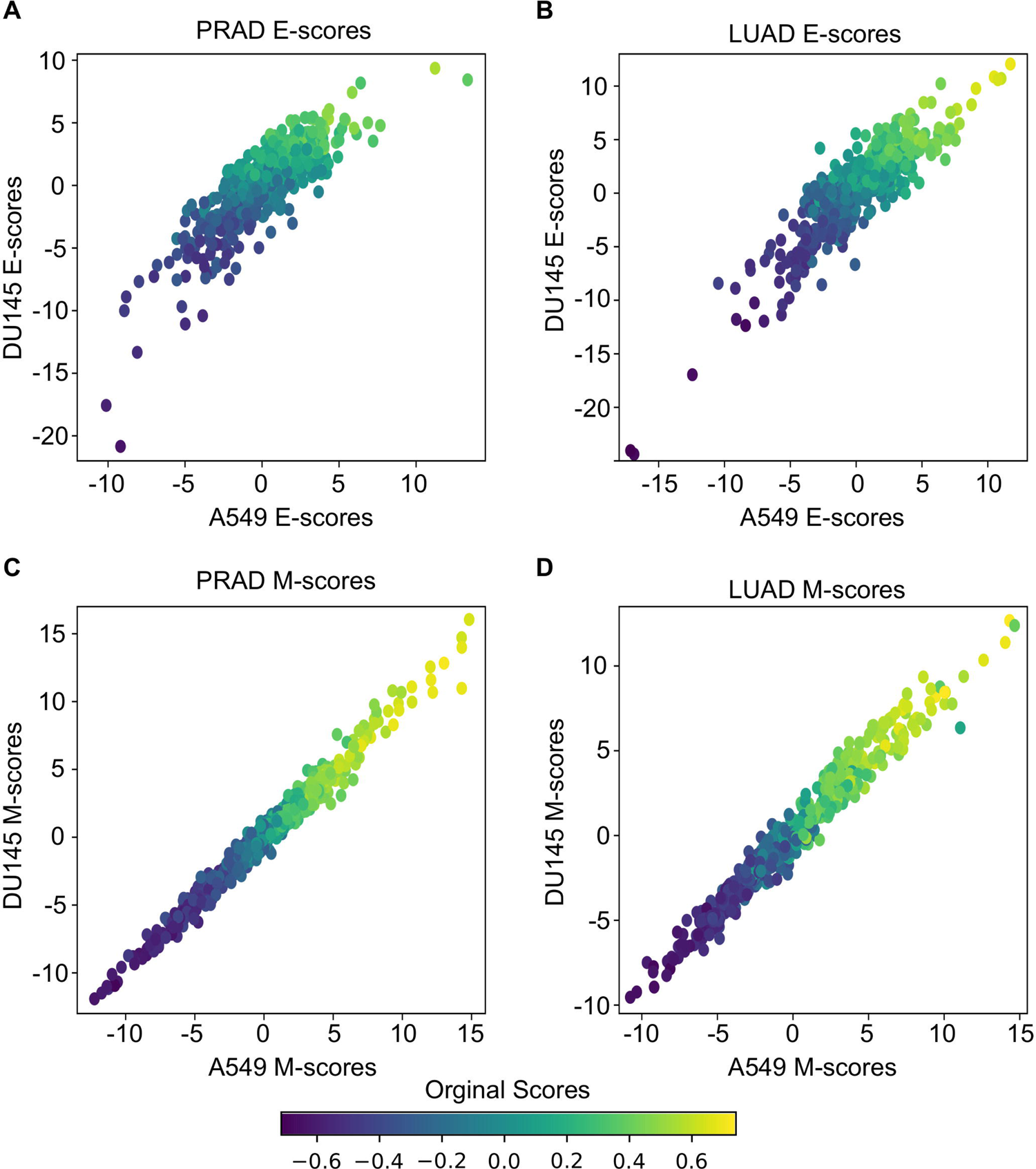
Transferring gsPCA models to TCGA data. (**A-B**) Scatter plots of E-scores for PRAD (A) and LUAD (B) from transferring gsPCA models built on A549 (x-axis) and DU145 (y-axis) data. The color of individual points indicates the original GSVA based E-score of the TCGA data set. (**C-D**) Scatter plots of M-scores for PRAD (C) and LUAD (D) from transferring gsPCA models built on A549 (x-axis) and DU145 (y-axis) data. The color of individual points indicates the original GSVA based M-score of the TCGA data set.

**Supplemental Figure S4.**
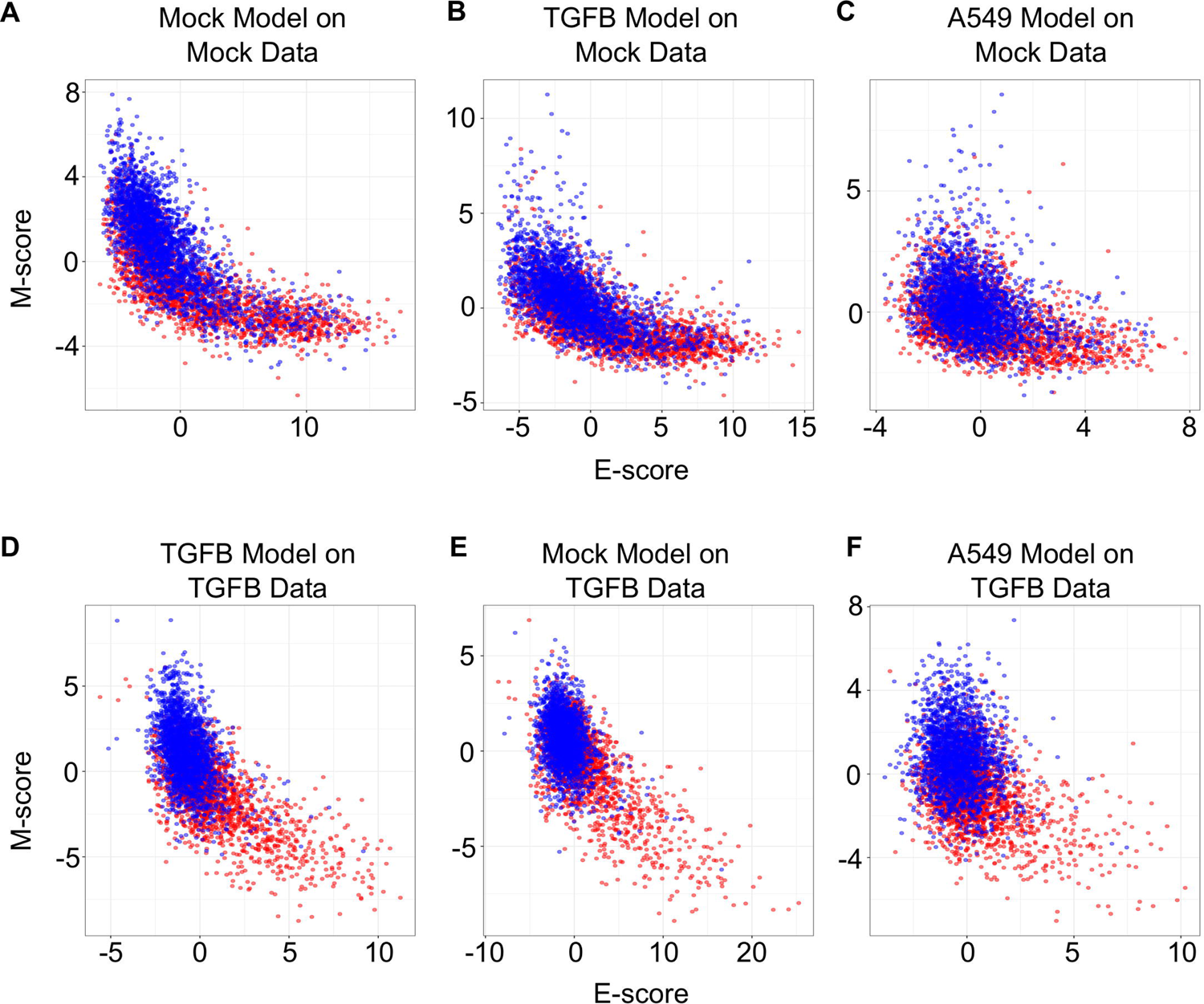
Transferring gsPCA models between temporal and spatial data sets. (**A-C**) Scatter plots of E (x-axis) and M (y-axis) scores for Mock spatial data from gsPCA models built on different data sets: Mock spatial data (A), TGF-β induced spatial data (B), and TGF-β induced A549 temporal data (C). The color of the sample indicates whether it originates from a cell in the inner-ring (non-motile, red) or the outer ring (motile, blue). (**D-F**) Scatter plots of E (x-axis) and M (y-axis) scores for TGF-β spatial data from gsPCA models built on different data sets: TGF-β induced spatial data (D), Mock spatial data (E), and TGF-β induced A549 temporal data (F). The color of the sample indicates whether it originates from a cell in the inner-ring (non-motile, red) or the outer ring (motile, blue).

**Supplemental Figure S5.**
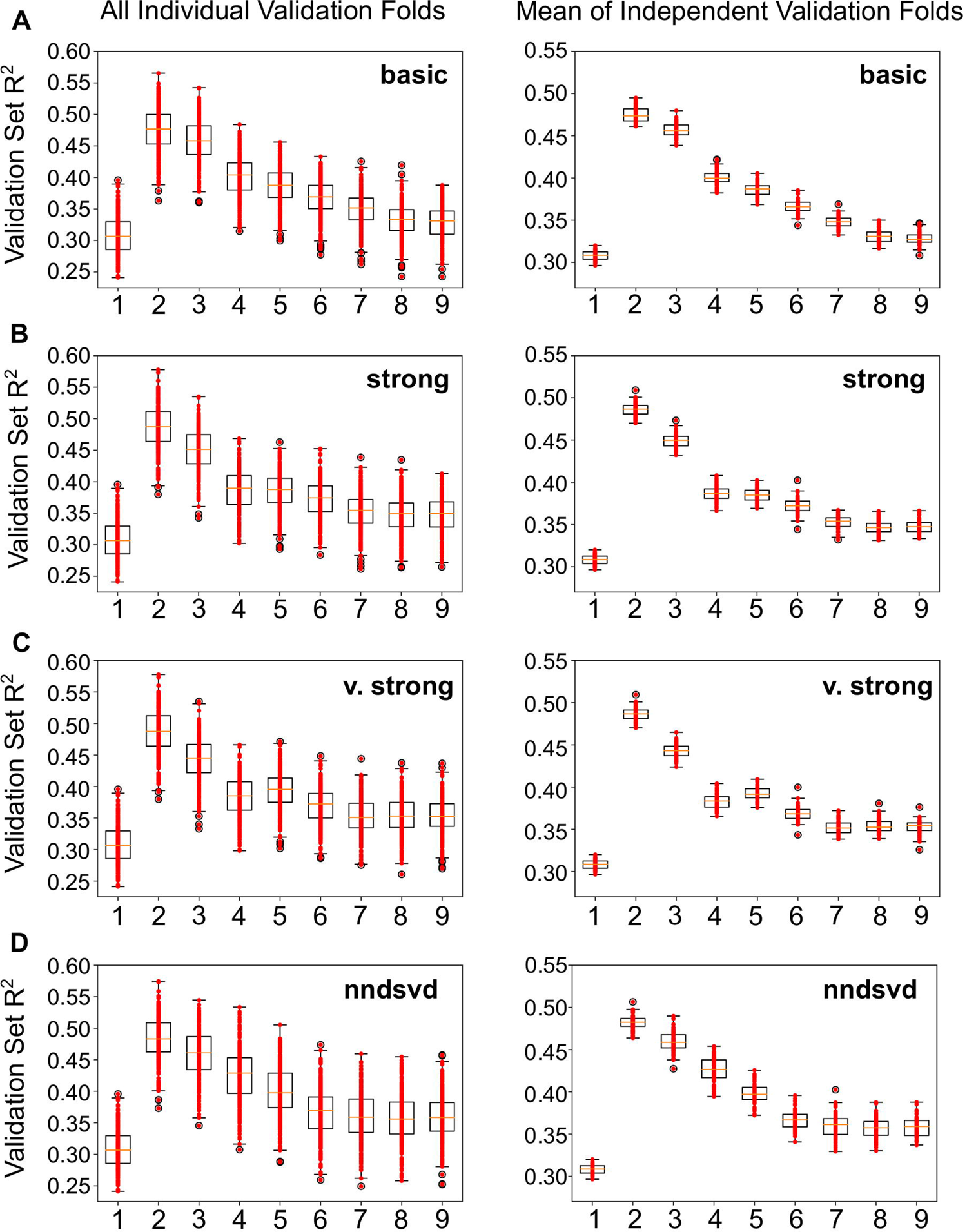
Performance of gsPCA models across cross-validation data sets. Boxplots showing Adjusted R^2^ of the linear model of gsNMF leading E and M dimensions across different number of model components (x-axis) and different convergence criteria: basic (algorithm standard, **A**), strong (1e-6 tolerance, 500 iterations, **B**), very strong (1e-9 tolerance, 2500 iterations, **C**), and nndsvd (initialization with non-negative singular value decomposition, 1e-6 tolerance, 500 iterations, **D**). Left and right panels are as in (**A**). The yellow line indicates the average of each distribution while the whiskers show 1.5 times the interquartile range. Red dots show the individual Adjusted R^2^ values in each distribution.

## Notes

### Competing Interest Statement

The authors have declared no competing interest.

### Summary of Updates

Added additional results section clarifying methodology and extending analysis of multiple gene sets to new data. Update pre-processing methodology for scRNA, including new data, and streamlined methods for clarity (extensive details on NMF/nnPCA can now be found in supplemental methods)

