## Supplemental Methods for "Interpretable, scalable, and transferrable functional projection of large-scale transcriptome data using constrained matrix decomposition"

**Background on non-negative PCA and NMF**

gsPCA uses the non-negative approach to PCA pioneered by Sigg and Buhnam (2008). In brief, the vectors of weights, *w*, used to define the first principal component of PCA is defined such that it maximizes the variance of the first component, i.e:

$${arg max}_{\boldsymbol{w}} w^{T}Cw$$

Where $C$ is the covariance matrix of the original data set $X$ and $w$ is unit vector (${||w||}^{2} = 1$)). In our case, $X$ is an $m$ by $n$ matrix of expression values where $m$ is the number of samples and $n$is the number of genes in the selected gene set. This method for determining $w$ can be treated as an expectation maximization problem where the original data is projected using the current estimate of $w$ ($y = Xw_{t}$) and this projection is used to re-estimate $w$ using the following minimization step:

$$w_{t+1}={arg min}_{w}\sum_{n=1}^{N} ||x_{n}-y_{n}w{||}_{2}^{2}$$

Where $x_{n}$ are the rows of the original data and $y_{n}$ are the rows of the projected data (1). This expectation-maximization formulation allows additional constraints on $w$, including forcing the component values to be non-negative. Note that the non-negativity constraint applies only to the weight components such that negative scores can still exist if there are negative values in underlying data, such as those produced by centering expression data to zero which we did for all gsPCA inputs. Subsequent components are calculated in the same way, under the constraint that they are orthogonal to the preceding ones.

NMF involves factorizing the original data matrix of non-negative values into two matrices whose product estimates the original data, i,e.:

$$X\tilde{=}WH$$

Where $X$ is the original matrix ($m$ by $n$)**,** $W$ is the transform matrix ($m$ by $p$), and $H$ is the component matrix ($p$ by $n$), such that $m$ is the number of rows in the original matrix (samples in our case), $n$ is the columns (genes in our case), and $p$ is the number of components used in the factorization. Because the original matrix is constrained to being non-negative, we subtracted the minimum of value of the scaled expression matrix from all values to create a non-negative input matrix. As a consequence, the values of the $W$ and $H$ matrices must likewise be non-negative such that product is non-negative.

**Cross-validation procedure**

As gsNMF involves an iterative minimization/maximization step, it can potentially yield variable results for same input data due to different initialization and convergence conditions as well as model parameters. To assess the performance of this approach, we implemented a cross-validation scheme using Scikit-learn. First, stratified k-fold sampling was used to split the A549 data set into six folds with equal proportions of samples from each time point. Each of the six-folds was reserved as a ‘testing’ set, while the remaining 5/6ths of the A549 data was used for training relative to that fold (i.e. six total data sets with exclusive training and testing data sets). The training data was further stratified into 5-folds, each of which was held out as a ‘validation’ data set to assess different model parameters which were trained on the remaining 4/6ths of the data. The initial 6-fold split was repeated 10 times, yield a total of 300 runs where models could be trained on 2/3rds of the A549 data set, validated on 1/6th and tested on 1/6th, with no repeated samples between the training, validation, and testing data, but roughly equal proportions of samples from different time points. Note that we used this procedure rather than standard 10-fold cross validation in order to ensure samples from all time points were well represented at each step of the process.

For assessment, we varied both the number of components used in the model (1 to 9) as well the convergence criteria: either the base setting of the algorithm (basic), a tolerance of 1e-6 and 500 iterations (strong), or tolerance of 1e-9 and 2500 iterations (very strong). We also used non-random initialization with non-negative SVD through Scikit-learn with the strong convergence criteria (nndsvd). The leading E and M dimensions for each model were selected using the correlation with EMT progression (most negative for E and most positive for M) and overall performance was assessed using the Adjusted-R^2^ of a linear model of E and M dimensions against time of EMT progression. For each set of parameters, we ran 10 random initializations over the training data and averaged Adjusted-R^2^ of the 10 models as applied to the validation data. We then considered two metrics of performance for each parameter set: the Adjusted-R^2^ across all validation folds individually and Adjusted-R^2^ averaged across each set of independent validation fold from the same training data set. Testing data was retained for evaluation of gsNMF against future approaches.

Overall, we found that a two component models gave the best Adjusted-R^2^ across all criteria (Figure S5). Across individual validation folds, we found that the mean Adjusted-R^2^ of two component models was significantly greater than the mean Adjusted-R^2^ of three components models across all convergence parameters (*p* < 0.05, Welch’s t-test). Among two component models, strong parameters (0.486) gave significantly better results than basic parameters (0.474; *p* = 0.0001, Welch’s t-test), but neither very strong parameters (0.486, *p* = 0.96) nor using non-random initialization (0.482, *p* = 0.99) significantly improved results. For the mean Adjusted-R^2^ of independent validation folds, we found the same pattern of significance across components and convergence parameters. Therefore, we chose a two component model as our baseline for gsNMF and used strong convergence parameters.

However, while these criteria perform well for our study, our choice is largely influenced by the type of analysis we perform. Therefore, for future applications of the approach, three main caveats should be considered. First, given our focus on dimension reduction and comparison of multiple gene sets on the same data or a single gene set across multiple data sets, we treat each gene set of having a single, primary relationship with the data set and seek the single best score to quantity that relationship and thus favor simpler models with fewer components. However, if the objective is to deconvolute multiple relationships from a larger set of functionally related or differential expressed genes, favoring more components would be reasonable. In such as case, it may be useful to refine each of the multiple components by selecting heavily weighted genes and generating new, smaller genes sets for each components. Additionally, assessment would be better done with an extended linear model or machine learning model using multiple-dimensions from each gene set as variables. Additionally, an alternative form of gsPCA using a cumulative algorithm that attempts to maximize the joint variation explained across all components (nscumcomp) might be useful for such an analysis. Second, with all of our data sets we were able to directly assess the performance of leading dimensions by assessing how well they explain a time variable or separate known groups, either using Adjusted R^2^ or the *f* probability. However, in cases where no such groups are known, an alternative criterion for choosing a leading dimension is required. In our study, the lead dimension from gsNMF tended to have the highest variance across random seeds (98.9% for E and 90.8% for M), but this is not constrained in the same as in gsPCA and the relationship tended to break down when using a large (>3) number of components. As such, for data without known groups to assess, we recommend gsPCA, which we have shown yields similar results to gsNMF overall for our data, but has the advantage of always generating a first PC which maximizes the variation explained regardless of the total number of components used. Finally, for applying NMF to the identification of samples and/or gene clusters on a full genome level, consider reviewing other works which more directly address the topics of low rank approximation and fitting very large number of components (2, 3), which are beyond the scope of our gene set focused approach.
